# Whole genome analysis identified a cefotaxime-resistant *Empedobacter brevis* GBW-1 isolate from ground beef encoding a novel metallo-beta-lactamase variant, *blaEBR-6*

**DOI:** 10.1101/2025.05.30.656955

**Authors:** Daniel Jones, Praful Aggarwal, Jamison Trewyn, Poojhaa Shanmugam, Kyle Leistikow, Troy Skwor

## Abstract

While investigating foodstuffs for ESBL-producing *Aeromonas* species on ampicillin dextrin agar with vancomycin and cefotaxime, a multidrug-resistant *Empedobacter brevis* strain GBW-1 was identified from ground beef. Phylogenetic analysis supports the interconnectedness of environment, humans and food driving this species evolutionary development. Antimicrobial susceptibility testing demonstrated resistance to gentamicin, carbapenems and third-generation cephalosporins. Whole genome sequencing of this strain detected a 3.74 Mb genome with 32.8% GC content containing 3,780 coding genes. Among these genes, at least three known antimicrobial resistance (AMR) genes were identified with *qacG, vanT* gene within the *vanG* cluster, and a novel variant of the metallo-β-lactamase *bla*_EBR-6_. This novel homologue, EBR-6, was compared against previously known EBR variants and was found to be closest to EBR-3 with an 84.98% amino acid identity match. Docking software predicted these mutations vary the binding to meropenem. Furthermore, nearly 100 annotated regions associated with mobile genetic elements, including the presence of three, separate *tra* operons were identified on the genome. Together, these findings implicate the importance of horizontal gene transfer in the acquisition of AMR among the emerging pathogen *Empedobacter brevis* and further stress its One Health nature.

**Importance:** As global trade and commerce continues to create a more interconnected world, the increasing frequency and spread of antimicrobial resistance amongst foodborne bacteria poses a significant challenge to public health. Here, we report the isolation of the multi-drug resistant *Empedobacter brevis* GBW-1, a bacterial strain harboring resistance against meropenem, third-generation cephalosporins, and gentamicin. Of particular note, this strain displays a novel variant of a metallo-β-lactamase (MBL) gene, *bla*_EBR-6_, as well as three distinct *tra* operons, suggesting enhanced capacity of horizontal gene transfer. These findings highlight how foodborne bacteria may serve as reservoirs and vectors for the further spread of resistance genes, reinforcing the necessity of utilizing a collaborative One Health approach to combat AMR across sectors.

## Background

Antimicrobial resistance (AMR) is one of the most challenging and urgent threats to global health. Microorganisms that exhibit multi-drug resistance (MDR) continue to be identified from a multitude of environments, underscoring the importance of utilizing a One Health approach to holistically and collaboratively tackle this ever-growing issue. Since 2017, The World Health Organization has compiled a list of drug-resistant priority pathogens threatening public health. In 2024, carbapenem- and third-generation cephalosporin-resistance among Enterobacterales remained in the ‘critical’ group of urgency (1). Due to their ubiquity within wide-ranging environments, capability of intra- and inter-species conjugation, and growing repertoire of antibiotic resistance genes (ARGs), the Gram-negative *Aeromonas* bacteria have been implicated as a possible indicator species for One Health-AMR surveillance (2, 3). However, the published literature investigating *Aeromonas-*AMR within food is limited. Our purpose was to help bridge this gap in knowledge by identifying ESBL-producing *Aeromonas* spp. in food. To identify *Aeromonas* spp., ground beef samples (25g) were diluted in 0.1% peptone, homogenized and plated on ampicillin dextrin agar with vancomycin (ADA-V; EPA Method 1605) and cefotaxime (1µg/ml). Plates were incubated at 30° C for 18 - 24 hours and presumptive cefotaxime-resistant *Aeromonas* isolates would appear as yellow colonies (Figure 1A). However, while confirmatory biochemical tests of one isolate revealed it is oxidase and indole positive, a negative trehalose result indicated it was not an *Aeromonas* isolate.

**Figure 1.**
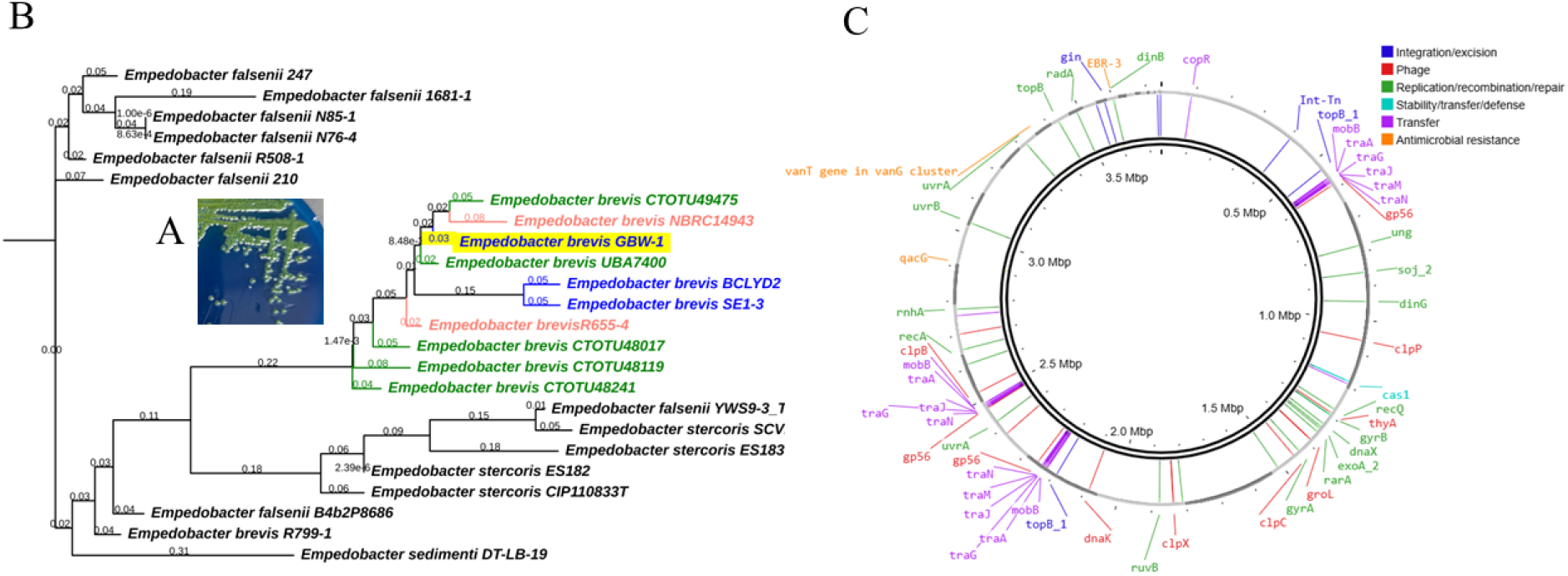
Isolation and whole genome analysis of *Empedobacter brevis* GBW-1. (A) *Empedobacter brevis* GBW-1 colonies on ADA-V agar subsequent to 24h incubation at 30° C. (B) Phylogenetic tree of *Empedobacter* genomes retrieved from NCBI. Strain GBW-1 (highlighted) is clustered with other *E. brevis* strains. Human isolates of *E. brevis* are in pink, while environmental isolates are in green and food isolates in blue. (C) Genetic map of *E. brevis* GBW-1 showing putative sites for ARGs along with the annotated protein families mediating the integration/excision, replication/recombination/repair, stability/transfer/defense or transfer of bacterial mobile genetic elements and phages.

Antimicrobial susceptibility using Kirby Baur disk diffusion assay on Mueller Hinton agar following CLSI standards (4) identified the isolate as resistant to gentamicin, amoxicillin/clavulanic acid, third-generation cephalosporins (i.e. cefotaxime and ceftazidime), and meropenem. The isolate was susceptible to fourth generation cefepime and monobactam, aztreonam. Due to the MDR nature of the isolate, genomic DNA was extracted using the Wizard Genomic DNA Purification Kit (Promega) and quantified via Qubit. Genomic libraries were prepared using the Illumina DNA Prep fragmentation kit and Illumina Unique Dual Indexes prior to whole genome sequencing on the Illumina NextSeq2000 platform (2×150bp paired reads).

Genomic assembly and annotation were submitted for the comprehensive genome analysis service at BV-BRC (5) and submitted to NCBI as Accession# JBLTIN000000000. Genome assembly was performed using Unicycler v0.4.8 and annotation was performed with RAST toolkit (RASTtk). This genome was found to be 3.74 Mb in size with 32.8% GC content.

Taxonomic classification was performed using Taxonomic database toolkit, GTDB-Tk v2.4.0 (6) and confirmed the isolate to belong to *Empedobacter brevis,* showing 98.18% average nucleotide identity and 84.9% alignment with the reference *E. brevis* genome (GCF_000382425.1).

Phylogenetic analysis (Figure 1B) using PhyloPhlAn v3.1.68 (7) further highlighted the genomic similarity of this strain, herein denoted as GBW-1, with other retrieved *Empedobacter brevis* isolates acquired from the environment (green), food (blue) and humans (pink). Formerly known as *Flavobacterium brevis* (8), *E. brevis* has been identified as an emerging pathogen associated with a variety of clinical infections in humans (9, 10), and is a common meat contaminant (11, 12) including among carbapenemase-producing organisms (13).

A total of 3,780 protein-coding genes (CDS), 65 transfer RNA (tRNA) genes, and 3 ribosomal RNA (rRNA) genes were predicted from *E. brevis* GBW-1. Among the predicted coding sequences (CDS), 3,591 had functional annotations, including 487 with Enzyme Commission (EC) numbers, 548 with Gene Ontology (GO) assignments, and 405 mapped to Kyoto Encyclopedia of Genes and Genomes (KEGG) pathways, while 261were hypothetical proteins. AMR genes were identified using the comprehensive antibiotic resistance database (CARD v6.0.3) (14). Only three AMR genes were identified: *vanTG* (vanT gene in vanG cluster), *qacG*, and a *bla*_EBR_ (Figure 1C). The vancomycin resistance observed by *E. brevis* GBW-1 on ADA-V is most likely due to the presence of *vanTG*. The presence of multidrug resistance efflux pump, *qacG*, has been associated with resistance to quaternary ammonium disinfectants (15). Lastly, BLAST analysis of the predicted amino acid sequence of *bla*_EBR_ exhibited only 84.98% similarity to EBR-3 as the closest related protein (Figure 2A), thus suggesting it as a novel member of the EBR metallo-β-lactamase family, EBR-6. From an evolutionary perspective, EBR-5 appears as the most recent descendent (Figure 2B) using the neighbor-joining method through MEGA11 (16). EBR-5 was identified in *Empedobacter stercoris* from chicken feces in China and exhibited decreased susceptibility compared to EBR-1 to cefotaxime and meropenem (17).

**Figure 2.**
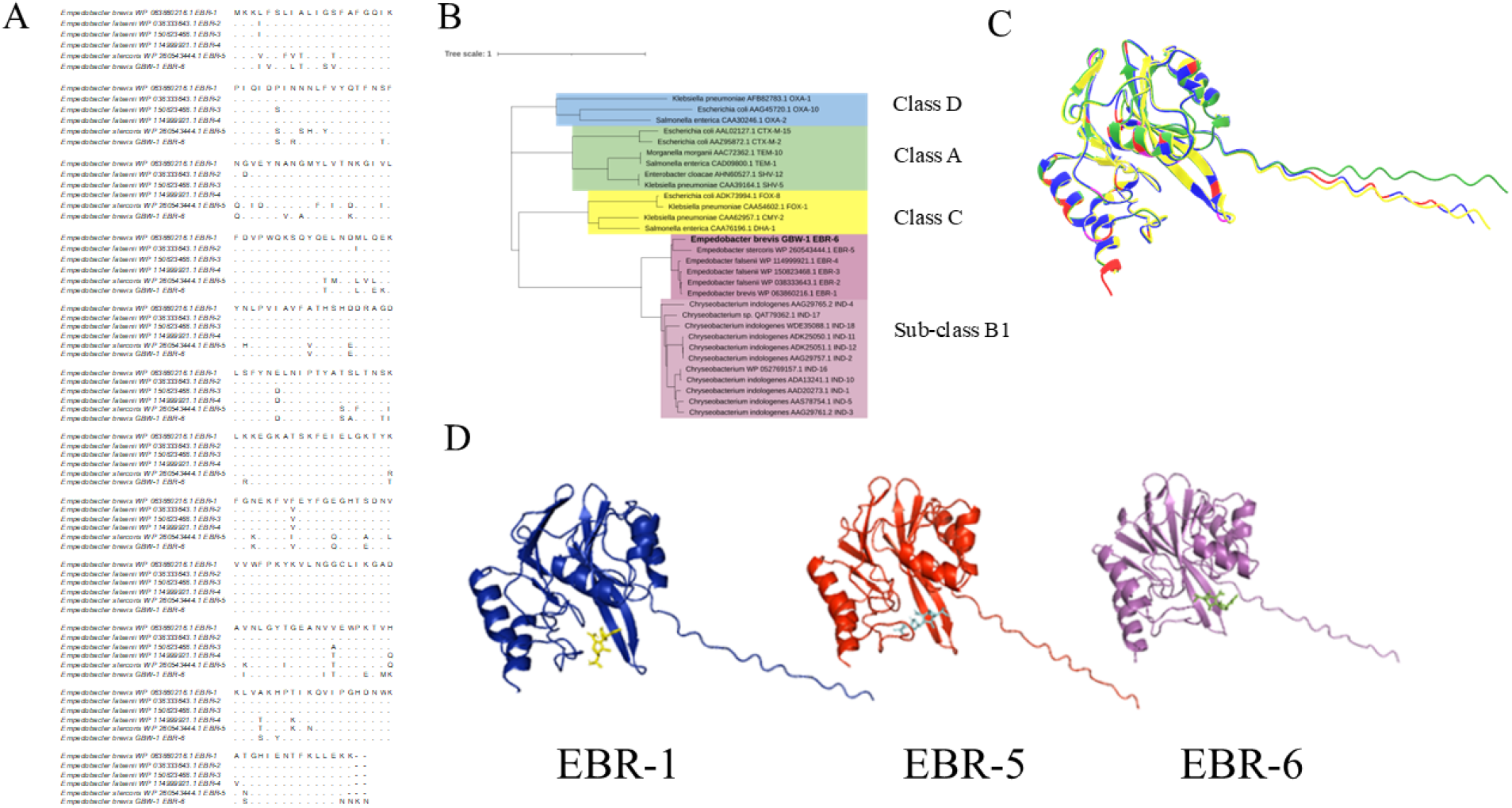
Phylogenetic analysis and variance of protein structure and meropenem docking among EBRs. (A) Amino acid comparison between all known EBRs. (B) Phylogenetic tree comparing amino acid sequences of EBR-6 to other beta-lactamases. (C) Comparison of predicted 3D protein structure of EBR-6 (blue) to EBR-1 (yellow) and EBR-5 (green). Am EBR-1 and -6 are in pink. (D) Computational predictions of meropenem docking between EBR-1, -5 and -6.

Due to the similarity to EBR-5, computational 3D renderings using AlphaFold3 (18) and ChimeraX 1.9 (19) were used to predict and compare EBR-1, EBR-5, and EBR-6 (Figure 2B). The 5 models constructed by AlphaFold3 were validated for quality using ProSA-web (Z-scores) (20). All the models have a Z-score of –8 and fall under the NMR region indicating significant quality (Suppl. Figure 1). The lowest Z-score EBR models and meropenem were cleaned, charged and hydrogenated for docking in BIOVIA Discovery Studio Visualizer (21) and AutoDock4 (22). Molecular docking was carried out using AutoDock Vina (23, 24). The best binding poses from each of the EBR-meropenem complexes were selected based on lowest binding affinity (kcal/mol), suggesting strongest predicted binding (Supp Table 1). The docking interactions were analyzed using BIOVIA to identify types of molecular interactions (hydrogen bonds or hydrophobic contacts) caused by the ligand (Suppl Figure 2). PyMOL (25) was used to estimate the differences in the binding of meropenem with each of the EBR proteins (Figure 2C).

Predicted common strong interactions (i.e. hydrogen, pi-sigma, pi-alkyl) with meropenem for the three EBR proteins were at Phe40, Tyr186, and His216 (Suppl Figure 3). Overall, EBR-5 and -6 both have more hydrogen and pi-bonds than EBR-1 (Suppl. Figure 3). Considering the decreased susceptibility of EBR-5 to third-generation cephalosporine, cefotaxime, and the carbapenem, meropenem, this suggests EBR-6 would have similar increased resistance as a carbapenemase, though experimental confirmation is required.

Lastly, putative mobile genetic elements were predicted using mobileOG-db v1.1.3 (26) and ran on the Proksee web server (27). Three separate *tra* operons were identified on *E. brevis* GBW-1, suggesting a capability to undergo horizontal gene transfer events (Suppl Table 2).

## Conclusion

Here we highlight the detection of a multi-drug resistant isolate of *E. brevis* from ground beef appearing as yellow on ADA-V agar supplemented with CTX, typically used for *Aeromonas* differentiation. WGS identified a novel metallo-β-lactamase, EBR-6, with multiple *tra* operons and numerous putative mobile genetic elements. EBR-6 exhibited similar docking patterns to EBR-5 suggesting decreased susceptibility to meropenem, though the computational nature of these genomic insights necessitates further experimental validation to confirm their functionality and mechanistic basis. Overall, due to the isolation of this strain from common foodstuffs, the isolation and identification of this EBR-6 gene in *E. brevis* further extends the evolutionary understanding of subclass B1 metallo-β-lactamases and implicates food as a major source of AMR. These data further emphasize the importance of reliable public health surveillance systems and the benefits of utilizing multisector approaches such as One Health in addressing the ever-growing issue of AMR.

